# Novel multiscale insights into the composite nature of water transport in roots

**DOI:** 10.1101/147314

**Authors:** Valentin Couvreur, Marc Faget, Guillaume Lobet, Mathieu Javaux, François Chaumont, Xavier Draye

**Affiliations:** Earth and Life Institute, Agronomy, Université catholique de Louvain, 1348 Louvain-la-Neuve, Belgium; Earth and Life Institute, Environment, Université catholique de Louvain, 1348 Louvain-la-Neuve, Belgium; Forschungszentrum Juelich GmbH, IBG3 Agrosphere, Juelich, Germany; Institut des Sciences de la Vie, Université catholique de Louvain, 1348 Louvain-la-Neuve, Belgium

**Keywords:** Aquaporins, Composite pathways, Cell hydraulics modelling, Root anatomy, Multiscale, Plasmodesmata, *Zea mays*

## Abstract

- MECHA is a novel mathematical model that computes the flow of water through the walls, membranes and plasmodesmata of each individual cell throughout complete root cross-sections, from a minimal set of cell level hydraulic properties and detailed root anatomical descriptions.
- Using the hydraulic anatomical framework of the *Zea mays* root reveals that hydraulic principles at the cell and root segment scales, derived independently by Katchalsky and Curran [1967] and Fiscus and Kramer [1975], are fully compatible, irrespective of apoplastic barriers leakiness.
- The hydraulic anatomy model accurately predicts empirical root radial permeability (*k*_r_) from relatively high cell wall hydraulic conductivity and low plasmodesmatal conductance reported in the literature.
- MECHA brings novel insights into contradictory interpretations of experiments from the literature by quantifying the impact of intercellular spaces, cortical cell permeability and plasmodesmata among others on root *k*_r_, and suggests new experiments efficiently addressing questions of root water relations.

**Symbols:**

*K*_PD_: single plasmodesma hydraulic conductance
*k*_r_: root radial hydraulic conductivity
*k*_w_: cell wall hydraulic conductivity
*L*_p_: cell plasma membrane hydraulic conductivity

## 1 Introduction

Vascular plant roots develop hydrophobic scaffolds (e.g., Casparian strips) diverting the course of water through cell membranes [Von Wangenheim et al., 2017], which filter as much as 60% of land precipitation on its way back to the atmosphere [Oki and Kanae, 2006]. In addition, root hydraulic and anatomical properties affect the plant water status, leaf growth [Caldeira et al., 2014] and crop water use under water deficit [Schoppach et al., 2014]. Yet, our quantitative understanding of these crucial hydraulic properties is hindered by the complex organization and regulation of sub-cellular elements such as aquaporins, which control the cell membrane permeability (*L*_P_) [Maurel and Chrispeels, 2001; Chaumont and Tyerman, 2014] and plasmodesmata, which connect adjacent cells protoplasts [Maule, 2008; Sevilem et al., 2013; Brunkard and Zambryski, 2017].

Due to the high hydraulic conductivity of cell walls, water pathways across root cortex are considered to be dominantly “apoplastic” [Steudle and Boyer, 1985]. In the vicinity of hydrophobic scaffolds, water passively crosses cell membranes through aquaporins and reach the symplasm. In cell layers where a hydrophobic suberin lamellae is embedded in the cell wall, such as the endodermis, the only way between neighbour cells is through plasmodesmata [Enstone and Peterson, 2005] which offer an uninterrupted “symplastic” pathway. Even though the resting diameter of plasmodesmata is approximately 50 10^−9^ m [Bell and Oparka, 2011], they are partly filled by a desmotubular structure, leaving water pathways that are only one order of magnitude wider than aquaporin aperture (2 to 3 10^−9^ m versus 2 10^−10^ m) [Terry and Robards, 1987; Tornroth-Horsefield et al., 2006].

A major difference between the principles of water flow within cell walls, through plasmodesmata and across membranes is the semi-permeability of the latter. Membranes exclude specific solutes, thus generating cell turgor from the same principle as the membrane osmometer [Katchalsky and Curran, 1967]. This semi-permeability is quantified as a “reflection coefficient” (***σ***, dimensionless) varying between 0 (fully permeable) and 1 (fully selective). In the first case, the water pressure difference (Δ*ψ*_p_, MPa) alone drives water flow, while in the second case the osmotic potential difference (Δ*ψ*_o_, MPa) contributes as additional driver. Such drivers generate water flow (*Q*, m^3^s^-1^) yet rate-limited by the media pore size distribution, summarized as its hydraulic conductance (*K*, m^3^MPa^-1^s^-1^):

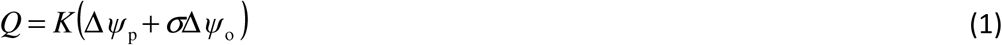

Fiscus and Kramer [1975] propose a similar principle that applies to the complex root structure. Their theory assimilates radial water transport to flow across a single semipermeable membrane:
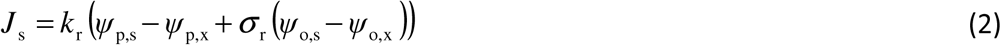

where *J*_s_ is the water flux at root surface (m^3^ m^-2^ s^-1^), *k*_r_ is the root radial conductivity (m^3^ m^-2^ s^-1^ MPa^-1^, note that here conductivity stands for conductance per surface area), *ψ*_p,s_ and *ψ*_p,x_ correspond to the water pressures at the root surface and in xylem vessels relative to atmospheric pressure (MPa), *σ*_r_ is the root reflection coefficient (dimensionless), and *ψ*_o,s_ and *ψ*_o,x_ correspond to the osmotic potential at root surface and xylem vessels, respectively (MPa).

Equation (2) is an effective equation as it merges the multiple cell-scale properties between root surface and xylem vessels into simple effective parameters *k*_r_ and *σ*_r_. Yet, whether or not this equation holds for any combination of hydraulic properties, anatomy and environment, is still unclear. In particular, systematically different *k*_r_ values were reported in hydrostatic and osmotic gradient experiments [Steudle and Frensch, 1989]. Such *k*_r_ differences were assumed to be the consequence of a substantial fully apoplastic pathway whose hydraulic conductivity would increase in hydrostatic pressure gradient experiments due to the water-filling of intercellular spaces [Steudle and Peterson, 1998].

The “root composite transport model” proposed by Steudle and Frensch [1989] considers that water follows pathways in parallel (apoplastic and cell-to-cell, the latter including symplastic and transcellular paths) that have different *k*_r_ and *σ*_r_ values. Root segment properties were linked to cell scale properties by integrating local conductivities and reflection coefficients using simplistic geometrical and hydraulic assumptions for each pathway [Steudle and Boyer, 1985]. Yet, the composite transport model does not constitute an explicit framework to investigate water pathways across individual cells, and for instance cannot explain why cortical cell *L*_p_ is not necessarily correlated to *k*_r_ [Hachez et al., 2012]. Furthermore, a validated root cross-section model may be used to target specific priority cell-level traits in a breeding context.

In this study, we develop a model of explicit root cross-section hydraulic anatomy (MECHA) built on the scientific community’s understanding of hydraulics at the cell scale, interconnected up to the root segment scale. One would expect that hydraulic theories at both scales should be compatible, though it was never explicitly verified. MECHA provides an explicit framework for testing such hypothesis, as it links quantitative hydraulic properties from the cell to the root segment scale. It also offers new insights to revisit conflicting interpretations of root water transport experimental results.

Firstly, we briefly introduce the model whose equations and parameterization are detailed in Notes S1 and section 5.1. The mathematical method to quantify water composite pathways across cell layers is detailed in Notes S3. Secondly, we check the compatibility of hydraulic theories at the cell and root segment scales. Thirdly, we assess the model predictions using literature data of maize *(Zea mays)* hydraulic properties at the cell and root segment scales. Eventually, we use MECHA to analyse and reinterpret previous statements concerning (1) the strong dependence of *k*_r_ to the water filling of cortical intercellular spaces [Steudle and Peterson, 1998], (2) the possible independence between *k*_r_ and cortical cell permeability [Hachez et al., 2012], and (3) the relative contribution of plasmodesmata to radial flow [Hukin et al., 2002].

## 2 Model

Root anatomical and cell hydraulic information were combined to build a finite difference model explicitly solving water pathways across individual cells of maize *(Zea mays)* roots.

The cell hydraulic information includes literature data of cell wall hydraulic conductivity (*k*_w_, which is an intrinsic conductivity in units of m^2^s^-1^MPa^-1^), cell plasma membrane permeability (*L*_p_, m s^-1^MPa^-1^), the hydraulic conductance of individual plasmodesmata (*K*_PD_, m^3^s^-1^MPa^-1^) and their frequency in each root tissue (see label “b” in Fig. 1). The cell geometry and connectivity information was collected from a cross-section image of the primary root of maize (Fig. 1a), vectorized with the program CellSet [Pound et al., 2012] (Fig. 1c). Together, they constituted the explicit root cross-section hydraulic anatomy (Fig. 1e). Further details of the parametrization are located in sections 5.1 and 5.2.

**Figure 1.**
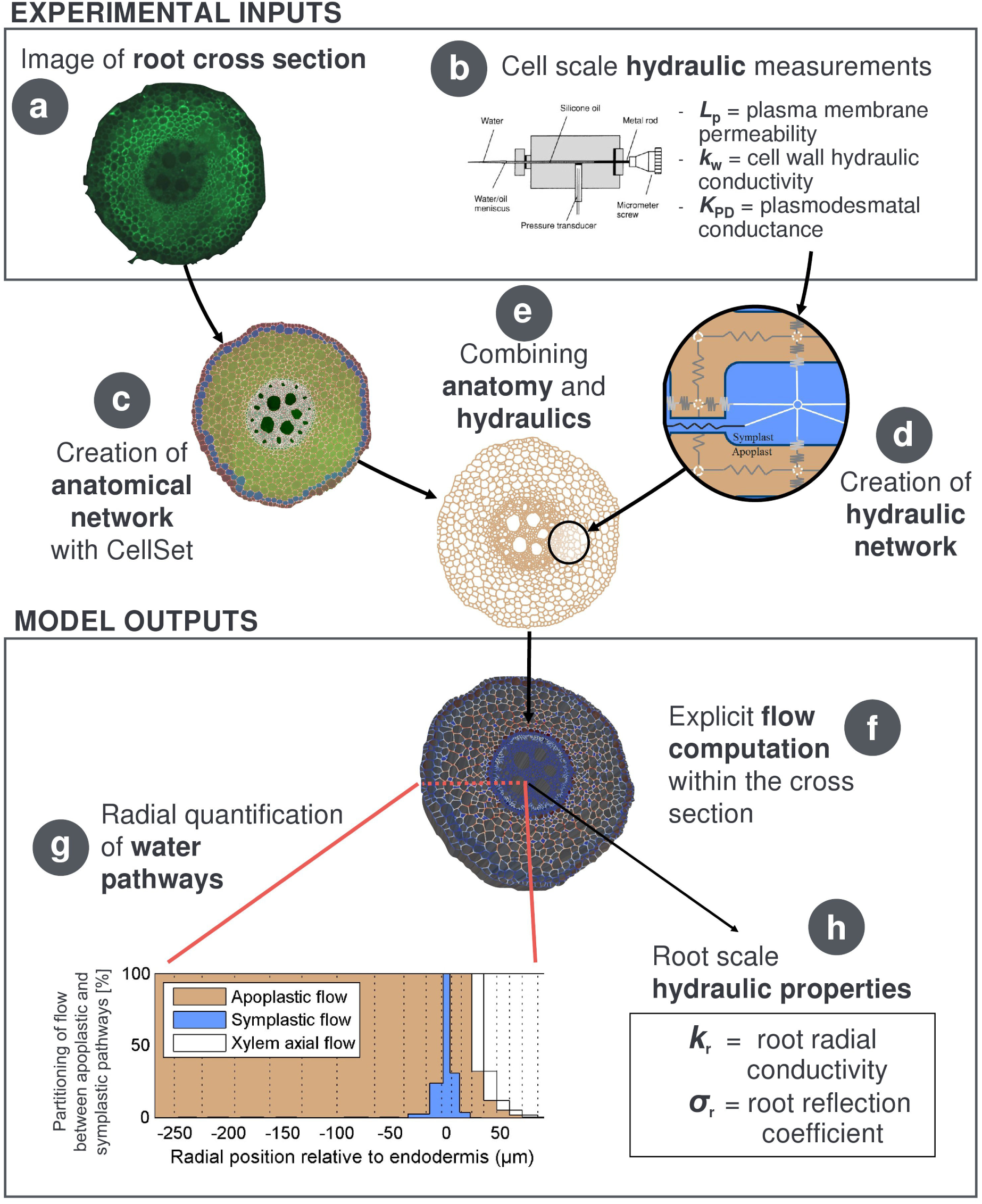
Overview scheme of the root cross-section hydraulic anatomy approach. A root cross-section microscope image (a) is treated through CellSet (c) in order to create the anatomical layout for cell-scale hydraulic principles (d) and associated properties (b), then forming the root hydraulic anatomy (e). The computed water flow rates across individual cells (f) also yield the root radial water pathways (g), root radial conductivity and reflection coefficient (h).

Computing water flow in the cross-section hydraulic anatomy yields flow rates across each individual hydraulic conductance (apoplastic, transmembrane and plasmodesmatal), as well as the distribution of water pressure in the apoplast and the symplast. Details on the water flow equations and solving method are given in Notes S1.

Examples of distributed water flow rates across cell walls, plasma membranes and plasmodesmata for relatively high *k*_w_ [Steudle and Boyer, 1985] and low *K*_PD_ [Bret-Harte and Silk, 1994] at various stages of apoplastic barrier deposition are described in section 5.1.

Furthermore, in Notes S3 we propose a mathematical method to quantify water radial pathways from local flow rates, which concentrates the essential flow information into a simpler graph (see panel “g” in Fig. 1).

## 3 Results

### 3.1 Hydraulic theories at the cell and root segment scales are compatible

Cell-scale hydraulic principles (Eq. (1)) and associated hydraulic properties (Tab. 2) were applied on a maize root cross-section anatomical layout in order to simulate water fluxes under various xylem and root surface boundary conditions (Tab. 3). Fig. 2a shows that the predicted radial fluxes ***J***_s_ at the root surface is proportional to the water potential difference between the root surface and xylem boundaries (*ψ*_p,s_ − *ψ*_p,x_ +*σ*_r_(*ψ*_o,s_ − *ψ*_o,x_)) as in the root segment scale theory of Fiscus and Kramer [1975] (Eq. (2)), with an R^2^ of 1.0000. The proportionality factor could thus be interpreted as equivalent to *k*_r_. Theories across scales therefore appear to be compatible, and this is valid for all combinations of cell scale hydraulic properties and apoplastic barriers leakiness.

**Figure 2.**
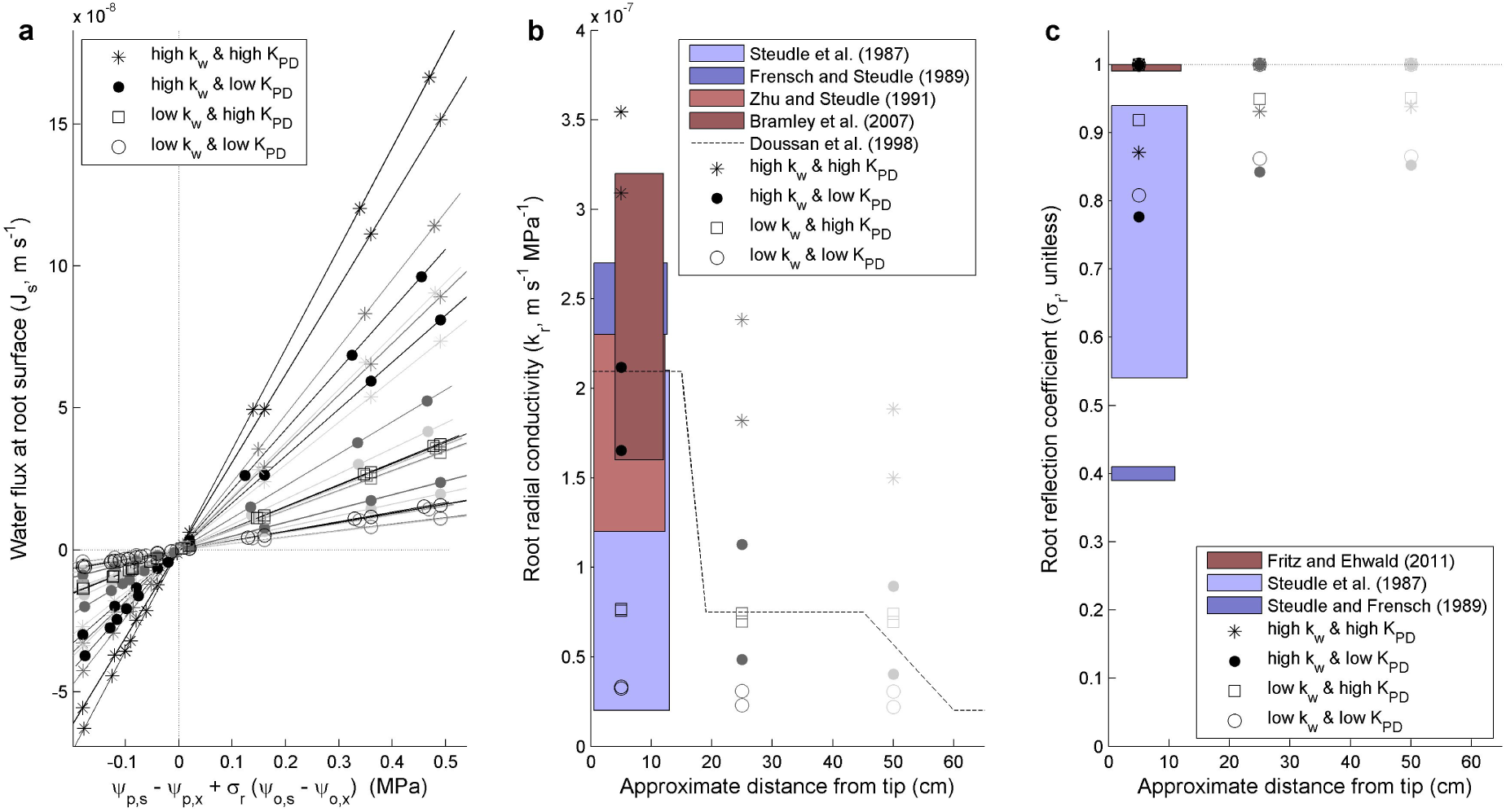
Cross-scale comparison of hydraulic principles and properties for various cell scale hydraulic properties (see legends) and levels of apoplastic barriers development (black: endodermal Casparian strip, dark grey: suberized endodermis with passage cells, light grey: suberized endodermis and exodermal Casparian strip). (a) Radial water fluxes simulated from cell-scale hydraulic principles in the hydraulic anatomy match the linear trend predicted from root segment scale principles for all tested boundary conditions (see Tab. 3) and hydraulic properties (see Tab. 2, with impermeable or leaky apoplastic barriers). (b) Comparison between empirical *k*_r_ ranges in maize primary roots (coloured rectangles and dashed line) and root segment *k*_r_ simulated from cell-scale hydraulic properties arranged in the maize hydraulic anatomy (symbols appear as couples, corresponding to the impermeable and leaky apoplastic barriers, the latter corresponding to the higher symbol). (c) Comparison between empirical mannitol *σ*_r_ ranges in maize principal roots (coloured rectangles) and root segment *σ*_r_ arising from cell-scale hydraulic properties arranged in the maize hydraulic anatomy (*σ*_r_ lower than 1 were obtained with the leaky apoplastic barrier).

Because it displays sensitivity to cell-scale hydraulic properties but not to boundary conditions, the upscaled *k*_r_ seems to be an intrinsic property of the hydraulic anatomy alone. This result suggests that the empirical observation of an impact of boundary conditions on *k*_r_ [Steudle and Frensch, 1989; Steudle and Peterson, 1998] are due to changes of cell-scale hydraulic properties (e.g., water-filling of intercellular spaces, or aquaporin gating quickly responding to the root osmotic environment [Hachez et al., 2012; Parent et al., 2009; Aroca et al., 2011]).

The simulations indicate that the additivity of pressure and osmotic potentials (after correction by the reflection coefficient *σ*_r_) is conserved from the cell to the root segment level. This additivity principle might also be preserved from root segment to whole plant scales, as suggested by experimental and modelling studies [Hamza and Aylmore, 1992; Schröder et al., 2014]. Like in Eq. (2), ***J***_s_ is not affected by alterations of cytosolic osmotic potential _(_*ψ*_o,c_, see Tab. 3), even though *ψ*_o,c_ was explicitly included in water flow equations across cell membranes (Eq. (S1)). This means that, when uniform, the effect of *ψ*_o,c_ somehow cancels out in the complete hydraulic network. Although radial gradients of root *ψ*_o,c_ were observed in transpiring maize plants [Rygol et al., 1993], they were not considered here. Their impact on water circulation inside roots will be investigated in a future study.

Models of water flow were not always found fully compatible across scales, as was the case in root water uptake models [Feddes et al., 1978; Couvreur et al., 2014; Javaux et al., 2013], or in hydrological models that rely on effective principles [Pokhrel and Gupta, 2010; Brynjarsdóttir and O’Hagan, 2014; Couvreur et al., 2016]. Lacking key processes or processes too complex to be represented with simple physical equations at either scale might explain an incompatibility. In this case, it seems that principles controlling water flow within roots were well captured, so that root segment hydraulics could be explained from underlying principles at the cell scale. Such compatibility opens avenues for the transfer of hydraulic information between cells and root segments.

### 3.2 Cell scale hydraulic properties injected in the maize hydraulic anatomy match empirical root segment *k*_r_

The radial hydraulic conductivity (*k*_r_) was frequently measured on the apical part of maize primary roots (up to 14 cm long) using a root pressure probe. Data from Steudle et al. [1987], Zhu and Steudle [1991], Frensch and Steudle [1989], and Bramley et al. [2007] cover one order of magnitude but mostly meet around 2 10^−7^ m s^-1^MPa^-1^ (see coloured areas in Fig. 2b). Root *k*_r_ data in mature root zones are rare, likely due to the presence of lateral roots that prevent independent measurement of the principal root segment *k*_r_ and complicate the attachment of the root to the probe. Doussan et al. [1998b] estimated a whole profile of root *k*_r_ along the maturation gradient by combining inverse modelling and cumulative flow experimental estimations from Varney and Canny [1993] (see dashed line in Fig. 2b, with development of secondary and tertiary endodermal walls between 15 and 19 cm from the tip, and exodermis development between 45 and 60 cm from the tip). MECHA parameterized with empirical cell-scale hydraulic properties yielded radial conductivities (see slopes in Fig. 2a, or symbols in Fig. 2b) that fell in the empirical range of maize root *k*_r_. Note that all symbols were paired, the higher and lower *k*_r_ respectively corresponding to leaky and impermeable apoplastic barriers.

The similarity of hydraulic properties measured and predicted across scales strengthens the validity of the hydraulic anatomy approach, particularly when considering that the reported *k*_r_ ranges for maize (overall 2 10^−8^ to 3 10^−7^ m s^-1^MPa^-1^, see boxes in Fig. 2b) appear as relatively narrow in view of the *k*_r_ values reported in the literature for other plants. For instance, McElrone et al. [2007] measured *k*_r_ values from 10^−5^ to 10^−4^ m s^-1^MPa^-1^ with an ultra low flowmeter in deep fine roots of live oak and gum bumelia, while Zarebanadkouki et al. [2016] obtained *k*_r_ values of 10^−6^ m s^-1^MPa^-1^ in undisturbed lupine primary roots, by combining deuterium tracing, inverse modelling and analytical root hydraulic functions developed by Meunier et al. [2017].

The simulated *k*_r_ profiles along the root maturity gradient (from black to light grey symbols) display a clear dependence on the cell wall conductivity (*k*_w_). The *k*_r_ profile simulated with the low *k*_w_ value proposed by Tyree [1968] (stars and circles in Fig. 2b) is indeed quite uniform compared to that obtained with higher *k*_w_ values measured by Steudle and Boyer [1985]. A higher *k*_w_ seems therefore more compatible with the important changes of root *k*_r_ frequently observed along the root of various plants and attributed to apoplastic barrier deposition [Sanderson, 1983; Doussan et al., 1998b; Barrowclough et al., 2000]. Such changes might have resulted from longitudinal alterations of cell hydraulic properties, but this is unlikely as cortical cell membrane conductivities were reported to be axially uniform beyond the elongation zone in maize and onion [Zimmermann et al., 2000; Barrowclough et al., 2000].

The combination of high cell *k*_w_ and low *K*_PD_ (Tab. 2) yields the best match with the empirical range of young maize root *k*_r_ and it reproduces qualitatively the axial trend estimated by Doussan et al. [1998b]. These properties generate a dominantly apoplastic water radial pathway in root cortex as compared to other combinations of cell properties (Fig. 3), which favour the symplastic (continuous blue stripes) and/or transcellular pathways (radial alternation of brown and blue stripes). If the cell-to-cell fraction of cortical radial water transport were to be as low as implied by high *k*_w_ properties, the plasma membrane aquaporins abundantly expressed in root cortex during the day [Hachez et al., 2006] would hardly participate to radial water flow across the cortex. Still, they may remain necessary for cell turgor regulation [Beauzamy et al., 2014; Chaumont and Tyerman, 2014]. The dynamic regulation of root *k*_r_ might then be specifically endorsed by those aquaporins highly expressed in the vicinity of apoplastic barriers, such as ZmPIP2;5 [Hachez et al., 2006].

**Figure 3.**
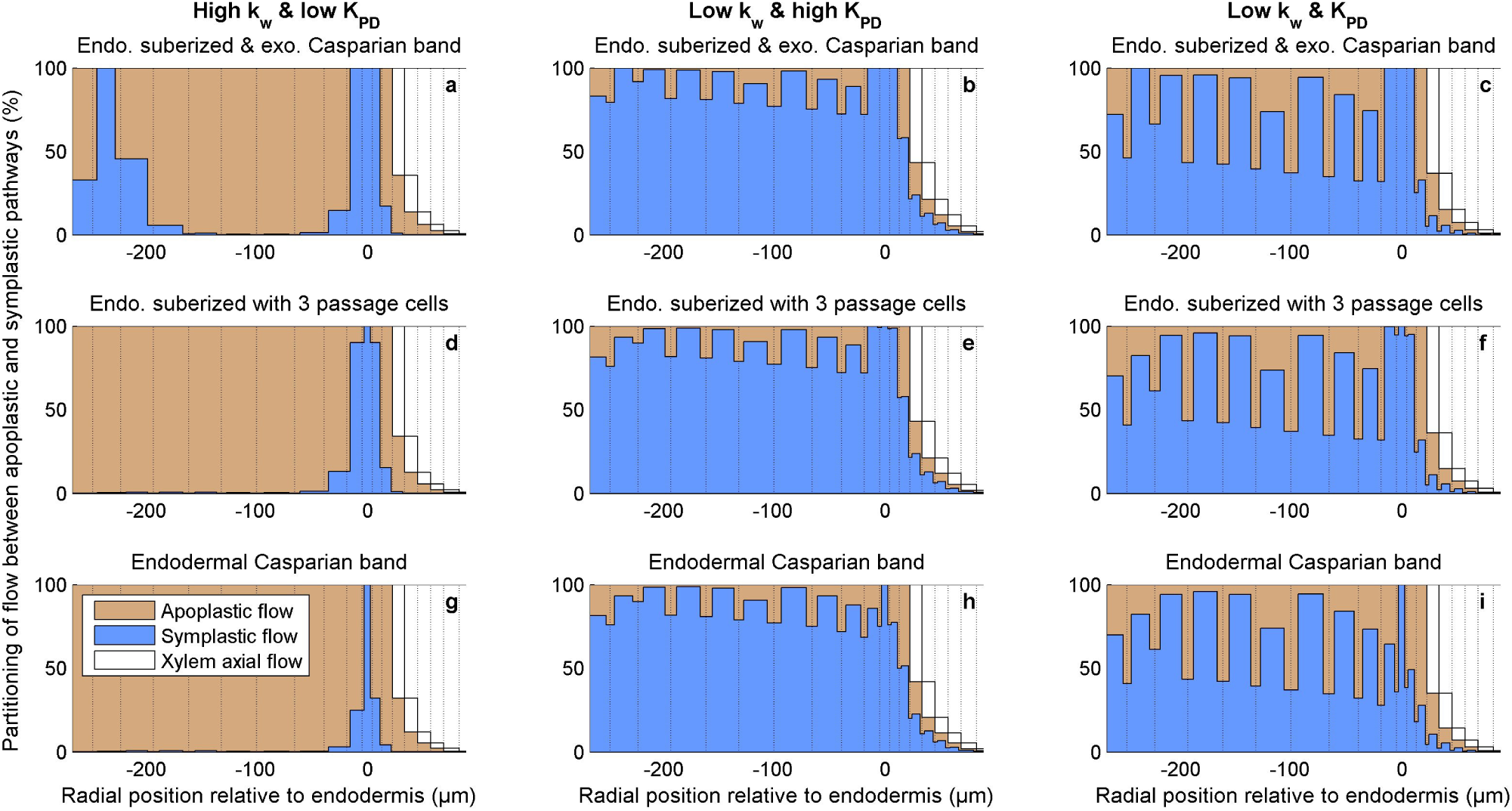
Radial partitioning of water pathways as percentage of the total radial flow, for the simple endodermal Casparian band stage (panels g, h, i), the suberized endodermis with 3 passage cells (panels d, e, f), and the fully suberized endodermis with an extra exodermal Casparian band (panels a, b, c). Impermeable apoplastic barriers and air-filled intercellular spaces were coupled to cell hydraulic properties from Tab. 2: high *k*_w_ and low *K*_PD_ (panels a, d, g), low *k*_w_ and high *K*_PD_ (panels b, e, h), low *k*_w_ and *K*_PD_ (panels c, f, i). Note that radial coordinates increase inwards.

The higher estimate of apoplastic barriers leakiness allows a significant fraction of fully apoplastic radial flow from root surface to xylem vessels (10.9 ± 5.6 %, for the 12 hydraulic scenarios reported at the end of section 5.1, not shown in Fig. 3 that has impermeable apoplastic barriers). Leakiness is also responsible for the reduction of *σ*_r_ below 1 (see Fig. 2c) even though all membranes reflection coefficients are equal to 1 (representative of the full selectivity of membranes to mannitol), as hypothesized in several studies [Steudle et al., 1987; Steudle and Frensch, 1989; Steudle and Peterson, 1998]. The experimental range of root *σ*_r_ for mannitol is wide (Fig. 2c) and even controversial as a few studies argue that values significantly lower than 1 stem from erroneous experimental interpretations [Knipfer and Fricke, 2010; Fritz and Ehwald, 2011]. The comparison between measured and upscaled *σ*_r_ is therefore tricky. In the upscaled model, *σ*_r_ appears to be an intrinsic property of the hydraulic anatomy and independent of boundary conditions, as *k*_r_.

### 3.3 Cortical intercellular spaces water-filling may not explain differences between osmotic and hydrostatic *k*_r_

Including water-filled intercellular spaces (see sections 5.2 and 5.5) increases the upscaled *k*_r_ by 2.9 ± 3.3 % on average with impermeable apoplastic barriers (maximum increase: 10%) and by 8.2 ± 5.1 % with leaky apoplastic barriers (maximum increase: 15%) in the 12 hydraulic scenarios. These values are by far lower than the typically observed ten-fold increase [Steudle and Frensch, 1989; Steudle et al., 1987] that Steudle and Peterson [1998] hypothesize to be the consequence of intercellular space water-filling. Despite their high hydraulic conductivity when water-filled, the impact of intercellular spaces on our upscaled *k*_r_ remains quite limited for the simple reason that they are in series with other hydraulic conductivities that limit the overall radial conductance.

The hydraulic anatomy approach therefore suggests that properties other than intercellular spaces water-filling likely generated the systematic differences between *k*_r_ measured in osmotic and hydrostatic gradient experiments. In line with this point, Knipfer and Fricke [2010] demonstrated that unstirred layers in the bathing medium are involved in the lowering of *k*_r_ in osmotic experiments in wheat, as no significant difference between *k*_r_ estimations in both types of experiments remain when circulating the root medium. Unstirred layers may provoke the underestimation or reduction of *k*_r_ in two ways: through the erroneous estimation of the osmotic potential difference between root surface and xylem |*ψ*_o,s_ − *ψ*_o,x_| in Eq. (2), or through alterations of cell hydraulic properties, such as *L*_p_.

A ten-fold change of *k*_r_ is quite substantial. From the empirical standpoint, closing maize aquaporins with H_2_O_2_ typically reduces *k*_r_ by less than four-fold (60% to 70% decrease [Ye and Steudle, 2006; Boursiac et al., 2008; Parent et al., 2009]). In MECHA, reproducing such alterations of cell hydraulic properties affects the upscaled *k*_r_ similarly in the youngest region of the root with high *k*_w_ and low *K*_PD_ (the most accurate scenario in Fig. 2b). Reducing *k*_AQP_ by 95% provoked a 56% to 72% decrease of the upscaled *k*_r_, for leaky and impermeable apoplastic barriers respectively. In another experiment, maize cortical cell *L*_p_ was reported to increase by 380% soon (< 2 hours) after the root medium water potential went from −0.075 to −0.34 MPa [Hachez et al., 2012]. Such alteration of *L*_P_ in all cells results in a 200 to 260% increase of the upscaled *k*_r_. Hence, our quantitative approach suggests that even though *L*_P_ is the most sensitive property controlling *k*_r_ before the endodermis suberization (see Fig. 4g), the reported empirical range of *L*_P_ changes alone is far from sufficient to generate ten-fold *k*_r_ alterations.

**Figure 4.**
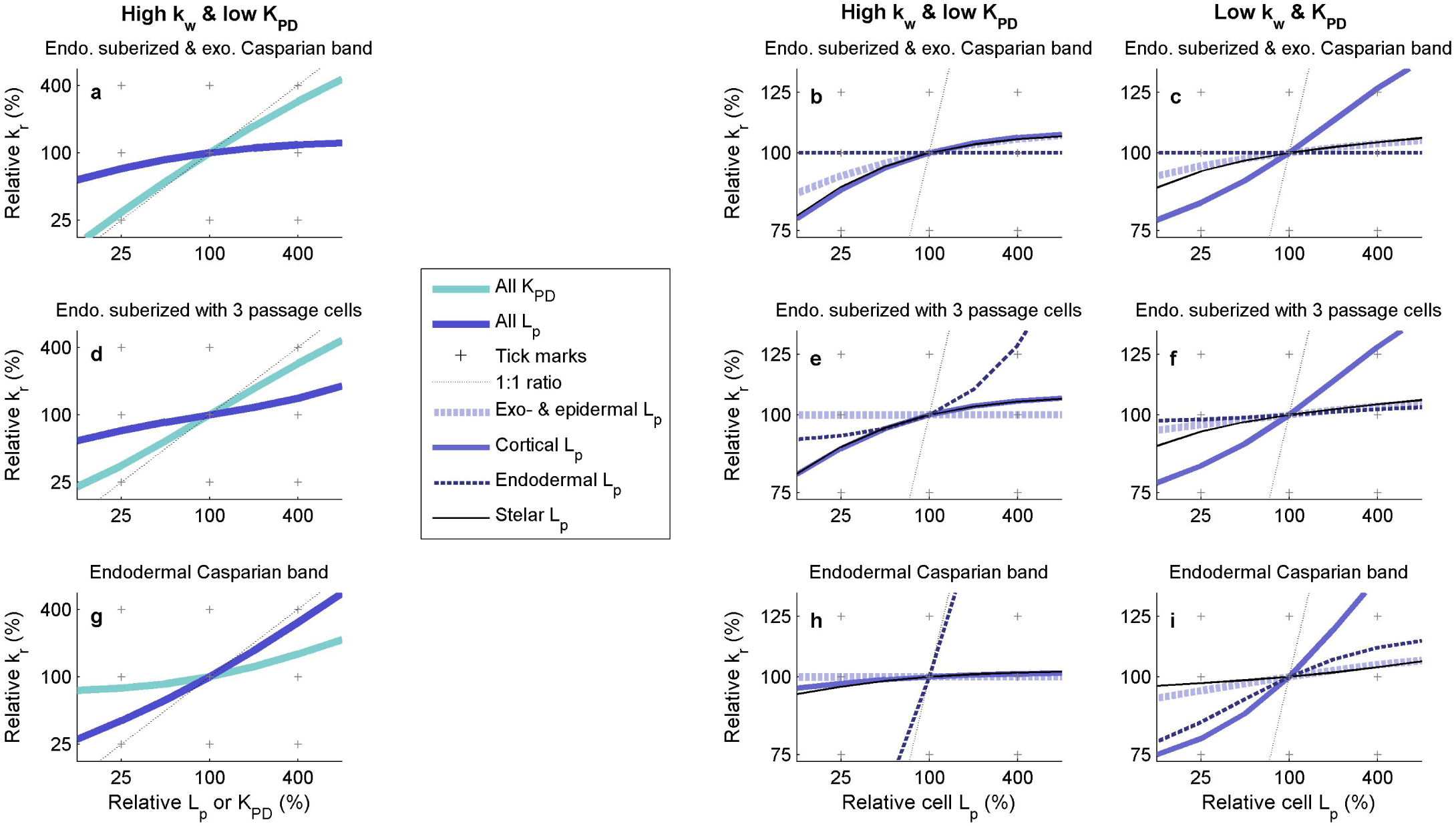
Variation of the cross-section radial hydraulic conductivity (ratio of the modified *k*_r_ to the default *k*_r_ as a function of the ratio of the modified cell hydraulic property to the default cell hydraulic property), for the simple endodermal Casparian band stage (panels g, h, i), the suberized endodermis with 3 passage cells (panels d, e, f), and the fully suberized endodermis with an extra exodermal Casparian band (panels a, b, c). In panels a, d and g, cell properties *(L*_p_ in blue or *K*_PD_ in cyan) were simultaneously altered in all tissue types, while in other panels, *L*_p_ was modified in specific tissue types in (epi- and exo-dermis in dashed light blue, cortex in solid blue, endodermis in dashed dark blue, and stelar cells in solid black). Panels b,e,h and c,f,i have a high and low cell wall hydraulic conductivities, respectiviely, and low *K*_PD_ (see Tab. 2). Note that a property with 100% sensitivity would imply null sensitivities of other properties, see Notes S4.

Our hypothesis is that a major part of reported *k*_r_ differences between hydraulic and osmotic experiments was an artefact stemming from the erroneous estimation of the osmotic driving pressure, as argued by Knipfer and Fricke [2010]. Plasmodesmatal aperture adjustment might also have contributed as it reportedly reacts to the root osmotic environment [Roberts and Oparka, 2003], but quantitative data is lacking so far.

### 3.4 Why maize root *k*_r_ correlates with the permeability of some cells and not of others?

Plasma membrane hydraulic conductivity (*L*_P_) appears to be the main property controlling *k*_r_ before the endodermal suberization in the scenario providing the best *k*_r_ prediction (high *k*_w_ and low *K*_PD_). However, with these hydraulic properties the endodermis is the only tissue whose *L*_P_ considerably affects the upscaled *k*_r_ (i.e. 69% of the overall sensitivity, versus 0.7% for cortical *L*_P_, see Fig. 4h). This result would explain the experimental observation that *L*_P_ and ZmPIP aquaporin expression in maize root cortex varied independently of *k*_r_ during osmotic stress [Hachez et al., 2012]. In MECHA, this feature occurs when the water stream bypasses cortical membranes (see Fig. 3g). The observed independence between cortical *L*_P_ and *k*_r_ cannot be reproduced with low cell wall hydraulic conductivities, which generate a high sensitivity of *k*_r_ to cortical *L*_P_ (up to 19% see Fig. 4i), and substantial cortical transmembrane flow (see Fig. 3h,i). Our model-assisted interpretations of experimental observations thus support the hypothesis of a higher range of cellulosic walls hydraulic conductivities from both angles of *k*_r_ sensitivity to apoplastic barrier deposition (section 3.2) and to cell *L*_P_ (section 3.4).

When the endodermis is partly suberized, the sensitivity to endodermal *L*_p_ remains higher than that to cortical *L*_p_ due to the transcellular water pathway across passage cells (Fig. 4e). It becomes negligible in more mature regions (see Fig. 4b) due to the isolation of cell plasma membranes from the apoplast following the endodermis full suberization.

Our results suggest that, before the suberization of the endodermis, the lack of correlation between cortical *L*_p_ and root *k*_r_ may simply stem from independent fluctuations of cortical *L*_p_ and endodermal *L*_p_, as only the latter significantly contributes to root *k*_r_. In order to verify it, we recommend to measure root *k*_r_, endodermal and cortical *L*_p_ before and during osmotic stress. Such experiment might also feed the question of whether separately regulated plasma membrane aquaporins fulfil different functions [Hachez et al., 2006], such as osmoregulation and radial permeability control.

### 3.5 Plasmodesmata may play a major role on root radial water transport despite their low conductance

Our results show that plasmodesmatal conductance might drastically alter water pathways from dominantly transcellular (Fig. 3c,f,i) to dominantly symplastic (Fig. 3b,e,h) when increasing *K*_PD_ by a factor 6, which would be achieved by expanding the plasmodesmatal inner spaces by a factor 2.4 only. Despite the relatively low plasmodesmatal conductance in the most accurate scenario (high *k*_w_ and low *K*_PD_), *k*_r_ sensitivity to *K*_PD_ is as high as 23% before endodermal suberization (Fig. 4g), and reaches 73% at maturity (Fig. 4a). When passage cells (occupying 4% of the endodermis surface in this cross-section) provide a bypass of plasmodesmata, the apoplastic flow converges toward the exposed plasma membranes of passage cells (see open arrowhead in Fig. S2c), contributing to 10% of water radial pathway (percentage visible in Fig. 3d). The remaining 90% still flows across the endodermis through plasmodesmata (see arrows in Fig. S2c) despite their low conductance.

The hypothesis of a major contribution of plasmodesmata to the water radial pathway has been formulated long ago [Clarkson et al., 1971], but was rejected by Hukin et al. [2002]. In the latter study, carboxyfluorescein was not radially translocated through plasmodesmata from the stele to the cortex in mature root segments (beyond the elongation zone). The authors thus concluded that plasmodesmata must be obstructed after the elongation zone and may not substantially participate to water transport either. However, calculations from Bret-Harte and Silk [1994] demonstrate that sugar transport across plasmodesmata massively relies on water flow, whose direction determines whether or not carboxyfluorescein can be translocated from the stele to the cortex. Considering that water flow across plasmodesmata may have been inwards after the elongation zone in the experiment of Hukin et al. [2002], our results indicating a major contribution of plasmodesmata to water transport remain compatible with the absence of carboxyfluorescein transport toward root cortex.

Advancing our understanding of water flow through plasmodesmata could involve the coupling of water tracing experiments (e.g., with deuterium [Zarebanadkouki et al., 2014]) to quantitative modelling of deuterium diffusion-convection in the root hydraulic anatomy. Each pathway would entail different deuterium distributions inside the root, and reproducing these distributions accurately in MECHA would take us one step further toward the verification of plasmodesmatal contribution to water radial flow across roots.

## 4 Discussion

MECHA is a novel hydraulic model which computes the flow of water through the walls, membranes and plasmodesmata of each individual cell throughout a complete root cross-section. From this fine scale, the model predicts root reflection coefficient (*σ*_r_) and radial permeability (*k*_r_). Hence it connects hydraulic theories across scales, based on detailed anatomical descriptions and experimental data on the permeability of cell walls (*k*_w_), membranes (*L*_p_), and plasmodesmata (*K*_PD_). Unlike Zhu and Steudle [1991] who define a single partition of water pathways (apoplastic versus cell-to-cell) for the overall trajectory between root surface and xylem, MECHA quantifies water composite pathways varying radially across successive cell layers, emphasizing the respective hydraulic function of each layer.

Using the hydraulic anatomical framework of a maize root as an example, MECHA reveals that hydraulic principles at the cell and root segment scales (Eqs. (1) and (2), respectively), derived independently by Katchalsky and Curran [1967] and Fiscus and Kramer [1975] are fully compatible, irrespective of apoplastic barriers leakiness. The upscaled maize root *k*_r_ matched the empirical range of measurements from the literature. In particular, the best reproduction of the *k*_r_ trend along the root maturation gradient was obtained using relatively high cell *k*_w_ [Steudle and Boyer, 1985] and low *K*_PD_ [Bret-Harte and Silk, 1994].

High *k*_w_ also yield root *k*_r_ that do not correlate with cortical cell *L*_P_ as observed experimentally [Hachez et al., 2012]. As the patterns of radial transmembrane flow and of *k*_r_ sensitivity to tissue *L*_P_ were comparable to the patterns of expression of specific aquaporins (ZmPIP2;5 [Hachez et al., 2006]), our results support the view that the function of specific aquaporins might be closely related to their localization. Regarding plasmodesmata, the model suggests that, despite their low conductance, they would play a major role in water radial transport across the suberized endodermis even in the presence of passage cells offering a direct transmembrane bypass, as argued by Clarkson et al. [1971].

The explicit hydraulic anatomy framework of MECHA offers the possibility to challenge the assumption of Steudle and Peterson [1998] that differences between osmotic and hydrostatic *k*_r_ stem from intercellular spaces water-filling. Our results do not support this hypothesis, as upscaled *k*_r_ were barely sensitive to such water-filling and independent of boundary conditions. Altered cell *L*_P_ coupled to inaccurate estimations of the osmotic driving pressure might explain the observation.

This enrichment of the composite transport model sheds new light on the field of plant water relations. Cutting-edge research on hydropatterning [Bao et al., 2014], hydrotropism [Dietrich et al., 2017], and apoplastic barriers [Barberon et al., 2016; Doblas et al., 2017] recently highlighted the need for a quantitative hydraulic framework to address questions related to pressure distribution and water flow direction at the cell scale. We expect MECHA to become a tool that will bridge the gap between protein regulatory pathways operating at the cell level and hydraulic behaviour at higher levels, pushing forward the potential of combined modelling and experimental approaches. Its compatibility with a root anatomical software [Pound et al., 2012] and functional-structural-plant-models [Javaux et al., 2008] has already opened avenues to investigate the relation between root architecture, anatomy and water availability [Passot et al., 2017].

The model comes with an online visualisation interface, available at <u>http://plantmodelling.shinyapps.io/mecha.</u>

## 5 Methods

### 5.1 Cell level hydraulic parametrization

The hydraulic conductance of individual plasmodesma (*K*_PD_) was evaluated from their geometry by Bret-Harte and Silk [1994] in maize, accounting for its partial occlusion by the desmotubule and increased viscosity of water in channels at the nanometre scale (3.4 mPa s). The obtained *K*_PD_ values range from 3.05 10^−19^ to 1.22 10^−18^ m^3^s^-1^MPa^-1^ (geometrical average: 6.1 10^−19^ m^3^s^-1^MPa^-1^, here referred to as “low *K*_PD_”). Their estimation is up to five orders of magnitude lower than previously calculated for barley roots by Clarkson et al. [1971] (10^−16^ to 10^−14^ m^3^s^-1^MPa^-1^) who neglected the presence of desmotubules, and an order of magnitude less than estimated experimentally by Ginsburg and Ginzburg [1970] in maize (3.54 10^−18^ m^3^s^-1^MPa^-1^, here referred to as “high *K*_PD_”). Note that in the latter case, a plasmodesmatal frequency of 0.48 μm^-2^ between the endodermis and the pericycle (as in Ma and Peterson [2001]) was assumed to turn the conductivity into a conductance per plasmodesma.

Single plasmodesmata conductances were assumed to be uniform (no data available on distributed plasmodesmata aperture) and scaled to the cell level through multiplications by plasmodesmata frequency (μm^-2^) and shared wall surface (μm^2^) of neighbouring cells. Plasmodesmatal frequency data in maize roots after cell elongation, within and between tissue types, were measured by Warmbrodt [1985] and Clarkson et al. [1987] (as reported by Ma and Peterson [2001], see Tab. 1). Shared wall surface estimations were based on the discretized root cross-section anatomy (see section 5.2: Root geometry).

**Table 1.** Plasmodesmatal frequencies within and at transitions between maize root tissue types. The missing frequency information was set at the average value by default.

| Tissue type, or transition between tissue types | Plasmodesmatal frequency ($\mu\text{m}^{-2}$) |
| --- | --- |
| Epidermis to exodermis | 0.54 |
| Exodermis to cortex | 1.14 |
| Cortex | 0.43 |
| Cortex to endodermis | 0.44 |
| Endodermis | 0.32 |
| Endodermis to pericycle | 0.48 |
| Pericycle | 0.35 |
| Pericycle to stelar parenchyma | 0.53 |
| Stelar parenchyma | 0.32 |
| Default | 0.4 |

Cell membrane hydraulic conductivity (i.e. conductance per membrane surface unit, associated to the minuscule *“k”* case) was measured with a pressure probe in maize root cortical cells by Ehlert et al. [2009]. As acid loading provokes the closure of aquaporins due to their protonation [Tournaire-Roux et al., 2003], the difference between hydraulic conductivities of control and acid load treatments (*k*_AQP_, 5.0 10^−7^ m s^-1^MPa^-1^) was attributed to aquaporins (the fraction of aquaporins that remained opened after the acid load was assumed negligible), while the remaining conductivity (2.7 10^−7^ m s^-1^MPa^-1^) includes the parallel contributions of the phospholipid bilayer (*k*_m_) and plasmodesmata (*k*_PD_). Note that the impact of plasmodesmatal conductivity cannot be distinguished from that of cell membrane conductivity with a cell pressure probe [Zhu and Steudle, 1991]. Hence, we used the geometrical-averaged *K*_PD_ with default plasmodesmatal frequency to obtain a default *k*_PD_ (2.44 10^−7^ m s^-1^MPa^-1^) and estimate the remaining *k*_m_ (2.6 10^−8^ m s^-1^MPa^-1^). For convenience, the sum of *k*_m_ and *k*_AQP_ was referred to as cell *L*_p_ in this paper.

Cell wall hydraulic conductivity (*k*_w_) was seldom quantified, but generally considered as least limiting the hydraulic media for water radial flow. Steudle and Boyer [1985] measured in soybean a value of 7.7 10^−8^ m^2^s^-1^MPa^-1^ (here referred to as “high *k*_w_”). They concluded that this relatively high value observed in hydrostatic pressure gradient experiments was likely due to the dominance of water flow in intercellular spaces, which was previously not accounted for. For instance, *k*_w_ was observed to be orders of magnitude lower in maize (2.5 10^−10^ to 6.1 10^−9^ m^2^s^-1^MPa^-1^ [Zhu and Steudle, 1991]) and isolated Nitella cell walls (1.4 10^−10^ m^2^s^-1^MPa^-1^ [Tyree, 1968], here referred to as “low *k*_w_”).

Casparian strips and suberin lamellae being hydrophobic, they were attributed null hydraulic conductivities, except in scenarios investigating the hypothesis of leaky apoplastic barriers. Lignified cell wall pores have the order of magnitude of 1 nanometre [Deng et al., 2016], which is small enough to fractionate between mannitol (C_6_H_14_O_6_) and ribitol (C_5_H_12_O_5_) [Fritz and Ehwald, 2011]. From Poiseuille law, we estimated that their intrinsic hydraulic conductivity could be no more than 10^−11^ m^2^s^-1^MPa^-1^, which was used as higher range for leaky apoplastic barriers. A summary of cell scale hydraulic properties is displayed in Tab. 2.

**Table 2.** Summary of cell scale hydraulic properties used in the analyses. Note that *L*_P_ variations were considered in the sensitivity analysis through variations around the default value. The hydraulic conductivity of cell walls in apoplastic barriers is either null or equal to 10^−11^ m^2^s^-1^MPa^-1^ when leaky.

|  | Values | Units | Sources |
| --- | --- | --- | --- |
| Default $L_p$ | $5.3 \cdot 10^{-7}$ | $\text{m s}^{-1} \text{ MPa}^{-1}$ | Estimated from Ehler et al. [2009]<br>and Bret-Harte and Silk [1994] |
| High $k_w$ | $7.7 \cdot 10^{-8}$ | $\text{m}^2 \text{ s}^{-1} \text{ MPa}^{-1}$ | Steudle and Boyer [1985] |
| Low $k_w$ | $1.4 \cdot 10^{-10}$ | $\text{m}^2 \text{ s}^{-1} \text{ MPa}^{-1}$ | Tyree [1968] |
| High $k_{w, \text{barrier}}$ | $1.0 \cdot 10^{-11}$ | $\text{m}^2 \text{ s}^{-1} \text{ MPa}^{-1}$ | Poiseuille law |
| Low $k_{w, \text{barrier}}$ | Zero | $\text{m}^2 \text{ s}^{-1} \text{ MPa}^{-1}$ | Full hydrophobicity |
| High $K_{PD}$ | $3.5 \cdot 10^{-18}$ | $\text{m}^3 \text{ s}^{-1} \text{ MPa}^{-1}$ | Ginsburg and Ginzburg [1970] |
| Low $K_{PD}$ | $6.1 \cdot 10^{-19}$ | $\text{m}^3 \text{ s}^{-1} \text{ MPa}^{-1}$ | Geometrical average of range from<br>Bret-Harte and Silk [1994] |

### 5.2 Root geometry

The hydraulic network geometry reproduced the anatomy of an aeroponically grown maize principal root, five centimeters from the tip (above the elongation zone, 0.9 mm diameter). A cross-section stained by immunocytochemistry for the cellular distribution of ZmPIP2;1/2;2, two plasma membrane aquaporins, pictured with an epifluorescent Leica DMR microscope (Wetzlar, Germany) as in Hachez et al. [2006], was segmented with the program CellSet [Pound et al., 2012]. Individual cells and structures were labelled manually (xylem, epidermis, exodermis, cortex, endodermis, pericycle and stelar parenchyma) based on their shape and position in the cross-section (Fig. 5a). Phloem elements and their companion cells could not be identified at that resolution. However, as they do not bear apoplastic barriers and represent a relatively small fraction of the cross-section, we assumed that their specific properties did not significantly impact root *k*_r_, and labelled them as their neighbour stelar parenchyma cells. Cortical intercellular spaces (204 on total in this crosssection) were recognized based on their relatively small size as compared to surrounding cells. They were attributed specific properties so that water could freely cross them when water-filled, while this path was blocked in scenarios with air-filled intercellular spaces (see blue polygons, i.e. null apoplastic fluxes, in Fig. 5b).

**Figure 5.**
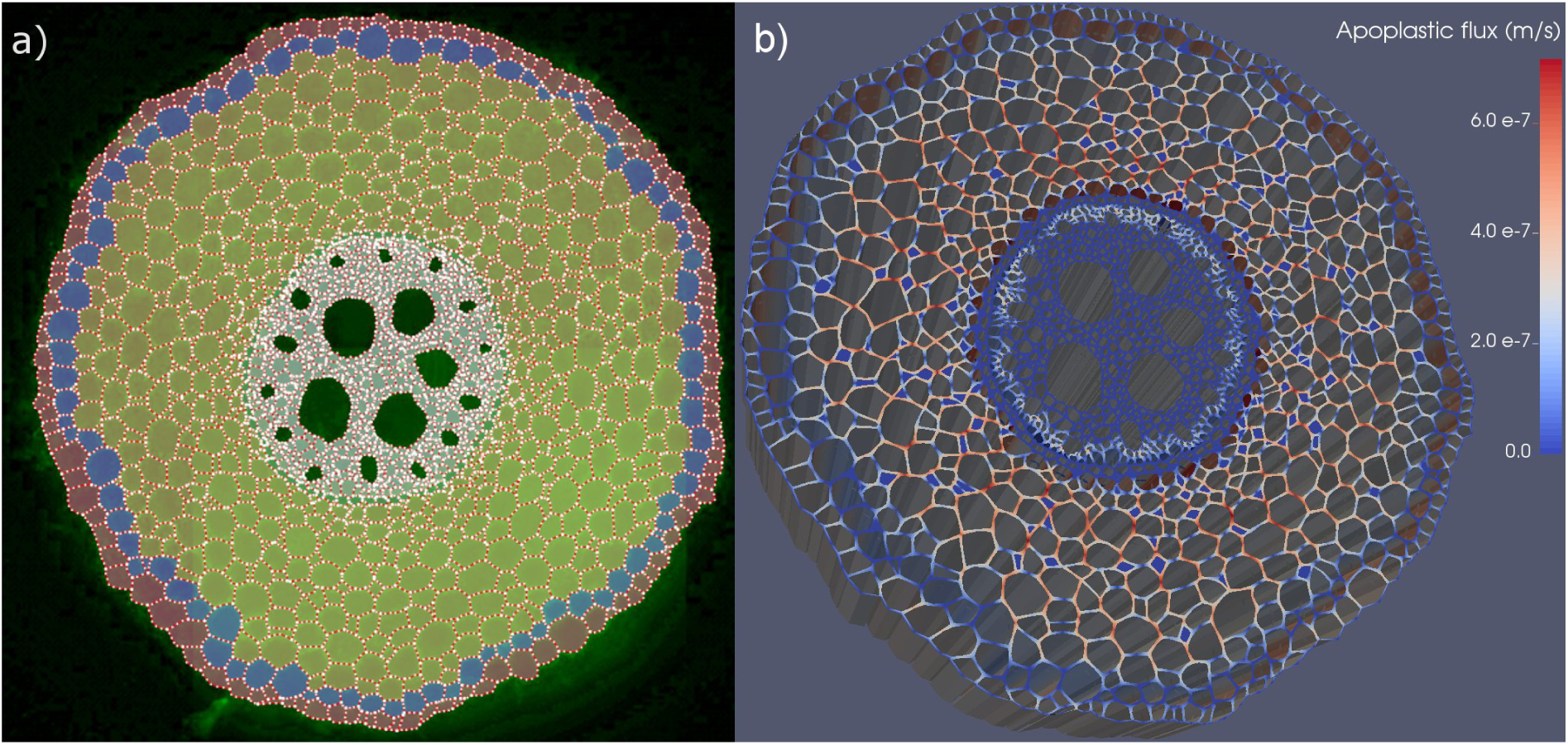
(a) Maize root cross-section with exodermis after segmentation in CellSet. Successive cellular tissue types from the periphery: epidermis, exodermis, cortex, endodermis, pericycle, other stele cells. Water potential boundary conditions were set at the epidermis surface and in xylem vessels (dark green). (b) Simulated water fluxes in cell walls, in m s^-1^, combining impermeable apoplastic barriers with high *k*_W_, low *K*_PD_ hydraulic properties and default boundary conditions (0.5 MPa water potential difference, generating an average uptake flux at root surface of 2.0 10^−8^ m s^-1^). Note that cell wall thickness is exaggerated to improve the visualization.

Labelled cells were grouped in layers, based on cell type (epidermis, exodermis, endodermis, and pericycle each constitute a cell layer) and cell connectivity. Cortical cells directly connected to the exodermis (through plasmodesmata) were assumed to belong to the first cortical cell layer. Similarly, cortical cells directly connected to the endodermis belonged to the last cortical cell layer. This process applied sequentially at each end of the not-yet-classified cortical cells. Classifying stelar cells was more straightforward as it started from the pericycle only. This layered classification is used in Fig. 3.

Three stages of apoplastic barrier development (structured deposition of hydrophobic material in cell walls) were selected for this study: (i) a simple endodermal Casparian band, (ii) an endodermis covered with suberin lamellae, except for three passage cells located in front of early metaxylem vessels, and (iii) a fully suberized endodermis (no passage cells) combined with an exodermal Casparian strip. In order to isolate the impact of apoplastic barrier deposition on root *k*_r_ and water pathways, the network geometry was conserved and cell wall hydraulic properties were adjusted at apoplastic barriers locations.

The cross-section was given a pseudo three-dimensional representation by attributing a height of 200 microns (typical length of maize elongated root cells) to the system. Transversal cell walls were thus included on top and at the bottom of the cross-section. In consequence, the top and bottom of cells were aligned in our model. As the mere presence of transversal walls affects *k*_r_ by less than 1.1% in all scenarios, we assumed that transversal wall alignment did not significantly change root hydraulic conductivity and water pathways, as compared to a more realistic non-aligned system. Note that only horizontal components of water flow were accounted for.

An example of distributed apoplastic water fluxes (i.e., flow rates per normal surface area) obtained from impermeable apoplastic barriers, high *k*_w_ and low *K*_PD_ hydraulic properties (Tab. 2) at the third stage of apoplastic barrier development (full endodermis and exodermal Casparian strip) is shown in Fig. 5b (other stages visible in Fig. S2). High velocities appear in red, mostly in the radial direction, corresponding to the direction of the water potential gradient. Low velocities in blue are mostly located at apoplastic barriers, where the apoplastic pathway is interrupted, and in tangential cell walls. While this detailed representation is appropriate to emphasize particular flow pathways in the vicinity of special features such as passage cells, the need for conciseness and clarity fostered the development of an alternative view of results: the radial representation of water pathways (see Notes S3, and Fig. 3).

### 5.3 Compatibility of hydraulic theories across scales

In order to verify the compatibility between hydraulic theories at the cell and root segment scales, we checked whether Eq. (2) holds when water flow across a root segment is simulated from cell scale hydraulic principles (Eq. (1)). If Eq. (2) holds in the aggregated system, J_s_ should scale linearly with the potential difference _“_*ψ*_p,s_−*ψ*_p,x_+*σ*_r_(*ψ*_o,s_−*ψ*_o,x_)”, for any given cell-scale hydraulic properties. The tested boundary conditions (Tab. 3) varied around experimental conditions reported in Steudle et al. [1987]: *ψ*_o,s_ the osmotic potential of the “Johnson-solution” (−0.02 MPa), *ψ*_p,x_ the equilibrated xylem pressure (+0.16 MPa), and *ψ*_o,x_ (−0.18 MPa) set to generate no flow when *ψ*_p,x_ equals +0.16 MPa. Cell cytosol osmotic potentials (*ψ*_o,c_) were set to the average value measured in maize cortex (−0.7 MPa) by Enns et al. [2000]. The verification was carried out for the combinations of cell hydraulic properties reported in Tab. 2, with both leaky and impermeable apoplastic barriers.

**Table 3.** Summary of boundary conditions selected for the tested scenarios.

| | $\psi_{p,s}$ (MPa) | $\psi_{o,s}$ (MPa) | $\psi_{p,x}$ (MPa) | $\psi_{o,x}$ (MPa) | $\psi_{o,c}$ (MPa) |
| --- | --- | --- | --- | --- | --- |
| Default boundary conditions | -0.01 | -0.02 | -0.34 | -0.18 | -0.7 |
| Various hydrostatic pressure gradients | -0.4 to 0.0,<br>+0.2 increment | -0.02 | -0.6 to +0.2,<br>+0.4 increment | -0.18 | -0.7 |
| Various osmotic pressure gradients | 0.0 | -0.2 to 0.0,<br>+0.1 increment | +0.16 | -0.18 | -0.7 |
| Various cytosol osmotic potentials | 0.0 | -0.02 | +0.24 | -0.18 | -0.6 to -0.8,<br>+0.1 increment |

Note that in this study, stelar cell walls osmotic potentials were assumed equal to that of xylem vessels (*ψ*_o,x_), other walls osmotic potentials were assumed equal to that of the root surface (*ψ*_o,s_), and membranes reflection coefficients were set to unity for all cells. A future study will focus on the higher levels of complexity implied by non-uniform reflection coefficients and osmotic potentials on water flow distribution in the root.

### 5.4 Compatibility of hydraulic properties across scales

An essential step strengthening the validity of the approach was carried out by verifying whether maize *k*_r_ empirical values from the literature match *k*_r_ predictions obtained using a typical root cross-section picture and empirical values of cell scale hydraulic parameters. This verification is novel as the present model is the first to have the ability to predict the root segment permeability from root section anatomy and cell hydraulic property distribution. The predicted cross-section radial hydraulic conductivity (*k*_r_, m s^-1^MPa^-1^) was calculated from the simulated root surface water fluxes (***J***_s_, m s^-1^) under default boundary conditions (see Tab. 3), using Eq. (2). Note that as argued by Fiscus and Kramer [1975] it would make sense to calculate *k*_r_ relative to the endodermis surface area. However, *k*_r_ empirical data reported in the literature is generally relative to the outer surface of the root. Hence, we complied with the latter approach.

### 5.5 Impact of intercellular spaces on root radial conductivity

In practice, hydraulic theories between cell and root segment scales may be compatible despite a non-linear relation between ***J***_s_ and boundary conditions, at the condition that *k*_r_ was altered by boundary conditions. For instance, Steudle and Peterson [1998] systematically measured maize *k*_r_ values that are one order of magnitude higher under hydrostatic than osmotic water potential gradients. Their interpretation is that in hydrostatic gradient experiments intercellular spaces get filled with water, thus increasing *k*_r_. Here we test the extent of *k*_r_ changes when intercellular spaces get filled with water in the model of the hydraulic anatomy, as compared with the air-filled case (i.e., infinite versus null conductivity of intercellular spaces). The analysis of *k*_r_ sensitivity to intercellular spaces water-filling was carried out for all cell hydraulic properties reported in Tab. 2, for both leaky and impermeable apoplastic barrier scenarios, the latter allowing a fraction of purely apoplastic flow, as hypothesized by Steudle and Peterson [1998].

### 5.6 Impact of cell plasmodesmata and plasma membranes permeabilities on root radial conductivity

Eventually we tested whether (and in which conditions) root *k*_r_ could be non-correlated to cortical cell membrane permeability as observed by Hachez et al. [2012], by running a sensitivity analysis of root *k*_r_ to various cell scale properties. The analysed properties were (i) cell membranes and plasmodesmatal hydraulic conductivities (for all cells simultaneously), and (ii) cell membrane conductivities (in specific tissues). This analysis was carried out for all cell hydraulic properties reported in Tab. 2, using the default boundary conditions from Tab. 3. Details of the sensitivity quantification were developed in Notes S4.

## Acknowledgements

All information about the model, including the source code, is available here: <u>https://mecharoot.github.io/</u>

A permanent version of the code used in the model is available here: ZENODO LINK

We also developed a web application to visualize typical outputs of our model. It was developed using the R Shiny framework and uses the following packages: ggplot2, ddply, reshape2. The web application is accessible here: <u>https://plantmodelling.shinyapps.io/mecha/</u>

MECHA was released under the GPL.2 open source licence.

We would like to thank Prof. C. Hachez for providing the root-cross section anatomical image. This work was supported by the Belgian National Fund for Scientific Research (FNRS), the Interuniversity Attraction Poles Programme-Belgian Science Policy (grant IAP7/29), and the “Communauté française de Belgique-Actions de Recherches Concertées” (grants ARC11/16-036 and ARC16/21-075).

V.C. and M.F. were supported by post-doctoral grants on the PAI MARS P7/29 project.

## Author contribution

V.C. and M.F. developed the code of MECHA. V.C., G.L., M.J., F.C., and X.D. gathered ideas and participated to the writing. G.L. developed the ShinyApp.

## Supporting information

### Notes S1: Detailed solution of water flow equations at the cell level

#### Overall description

The topology of the hydraulic network comprised symplastic and apoplastic “nodes”, where water potentials were defined. Nodes were connected by hydraulic conductances conveying water (Fig. S1). As we focused on steady flow conditions, which happens when root water uptake reaches a stable rate, water contents of components of the system remain constant (even though water crosses them). Hence, the network needed no capacitance. Furthermore, the hydraulic resistance of the cytosolic compartment was assumed negligible as compared to membranes. Since water could freely bypass organelles, a single symplastic node was located at the centre of each protoplast. The apoplast was discretized in apoplastic blocks, according to cell wall geometry. Apoplastic nodes were located at the centre of each apoplastic block and at their junctions (Fig. S1).

Water was assumed to flow through three types of hydraulic media characterized by specific conductivities (Fig. S1): transmembrane hydraulic conductances (in light grey), cell wall hydraulic conductances (in dark grey), and plasmodesmata hydraulic conductances (in black), see parametrization in section 5.1. As shown in Fig. S1, nodes at junctions between apoplastic blocks were not directly connected to symplastic nodes; their role was to traffic water in the apoplast. Note that only two plasmodesmata conductances were displayed for readability concerns, but each protoplast shared such conductances with all neighbouring protoplasts.

Three principles were used to compute water flow in the multicellular hydraulic network. Firstly, equations analogical to Ohm’s law governed water flow through individual hydraulic conductances (i.e. the flow rate equals the product of the hydraulic conductance by the potential difference), following Katchalsky and Curran [1967] (see Eq. (1)). As cell walls and plasmodesmata are not selective for most solutes, their reflection coefficients were set to zero. For a solute like mannitol, the osmotic potential difference would intervene fully (cell membrane *σ* equal to 1 [Zhu and Steudle, 1991]). Secondly, equations analogical to Kirchhoff’s law ensured mass conservation (i.e. each node’s water input equals its water output under steady flow conditions). Thirdly, water pressure boundary conditions were set at root outer surface and in xylem vessels, while osmotic potentials were defined for all nodes (see selected boundary conditions in Tab. 3).

#### Flow equations

For each type of cell level conductance, geometrical aspects were accounted for in order to turn an intrinsic conductivity into a conductance between two nodes. Water flow rates (i) between cell wall and neighbouring symplastic nodes across a portion of cell membrane, (ii) between a cell wall node and a neighbouring junction node, and (iii) between neighbouring intra cellular nodes across plasmodesmata (resp. *Q*_TM_, *Q*_apo_, and *Q*_PD_, m^3^s^-1^) were calculated as follows:
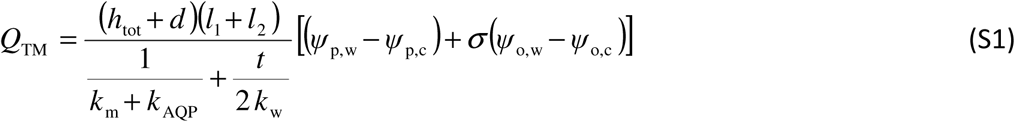

where *h*_tot_ (m) is the height of the cell in the root axial direction (here approximately 2 10^−4^ m when removing the cell wall thickness), *d* (m) is the horizontal distance between the cell wall and the cell centre, *l*_1_ and *l*_2_ (m) are horizontal distances between the cell wall node and neighbouring junctions nodes, *t* (m) is the wall thickness between two neighbouring cells (here 1.5 microns), *k*_m_(m s^-1^MPa^-1^) is the contribution of the phospholipid bilayer to the plasma membrane hydraulic conductivity, *k*_AQP_ (m s^-1^MPa^-1^) is the contribution of aquaporins to the plasma membrane hydraulic conductivity, *k*_w_ (m^2^s^-1^MPa^-1^) is the intrinsic cell wall hydraulic conductivity, *ψ*_p,w_ (MPa) is the cell wall pressure potential, *ψ*_p,c_ (MPa) is the cytosol pressure potential, *σ* (−) is the cell plasma membrane reflection coefficient, *ψ*_o,w_ (MPa) is the cell wall osmotic potential, and *ψ*_o,c_ (MPa) is the cytosol osmotic potential. Note that the factor *d* intervenes in order to approximate the contribution of horizontal portions of the plasma membrane at the top and bottom of the cell.
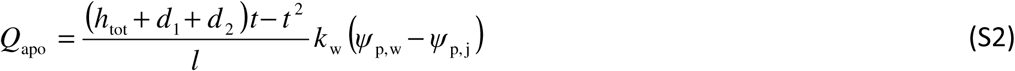

where *d*_1_ and *d*_2_ (m) are horizontal distances between the cell wall portion and neighbouring cell centres, *l* (m) is the horizontal distance between the cell wall node and the neighbouring junctions node, and *ψ*_p,j_ (MPa) is the cell wall junction pressure potential.
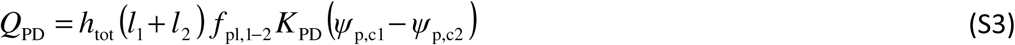

where *f*_pl,1-2_ (m^-2^) is the plasmodesmatal frequency between the neighbouring cells 1 and 2, *K*_PD_ (m^3^s^-1^MPa^-1^) is the hydraulic conductance of a single plasmodesma, *ψ*_p,c1_ and *ψ*_p,c1_ (MPa) are intracellular pressure potentials of neighbouring cells 1 and 2.

An equation analogical to Kirchhoff’s law governs mass conservation at each node in the network:
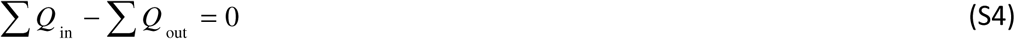

where *Q*_in_ refers to fluxes (*Q*_TM_, *Q*_apo_, and/or *Q*_PD)_ defined positive toward the node (its water potential is on the right in the potential differences), and *Q*_out_ stands for fluxes defined negative toward the node. Note that the right-hand-side term is null because in stationary flow conditions there is no accumulation of water possible at any location in the network.

#### Solving method

Let N_u_ be the number of nodes with unknown pressure potential, N_BC_ the number of boundary condition nodes (with known pressure potential), and N_tot_ the total number of nodes. A matrix form of the equation system can be constructed as in Doussan et al. [1998a]. The unknowns are grouped in the vector of pressure potentials ψ_p,u_ (N_u_ × 1, MPa), which is solved by left-multiplying both sides of the following equation by the inverse of the matrix of connected conductances in the domain of unknown pressure potentials C_p,u_ (N_u_ x N_u_, m^3^s^-1^MPa^-1^):
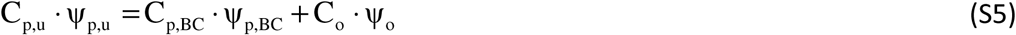

where C_p,BC_ (N_u_ × N_BC_, m^3^s^-1^ MPa^-1^) is the matrix of connected conductances in the domain of known pressure potentials (soil and xylem boundary conditions), ψ_p,BC_ (N_BC_ × 1, MPa) is the vector of known pressure potentials, C_o_ (N_u_ × N_tot_, m^3^s^-1^MPa^-1^) is the matrix of connected membrane conductances multiplied by their reflection coefficient, ψ_o_ (N_tot_ × 1, MPa) is the vector of osmotic potentials.

C_p,u_ and C_p,BC_ matrices are built as follows: for each node “i” with unknown pressure potential (index ranging between 1 and N_u_), the conductance to the connected node “j” with unknown pressure potential is added up to C_p,u_(i,i) and subtracted from C_p,u_ (i,j). The conductance to the connected node “k” with known pressure potential (boundary condition node, with index ranging between 1 and N_BC_) is added up to C_p,u_(i,i) and C_p,BC_(i,k). Besides, the C_o_ matrix is built as follows. For each node “i” with unknown pressure potential (its index within N_tot_ is called “i_tot_”, and differs from that within N_u_), the conductance to the membrane-connected node “j_tot_” multiplied by its reflection coefficient is added up to C_o_(i,i_tot_) and subtracted from C_o_(I,j_tot_).

After solving the system for pressure potentials, water fluxes are solved with the previously displayed equations of *Q*_TM_, *Q*_apo_, and *Q*_PD_.

### Notes S2: Visualization of cell-scale composite water flow patterns at different maturity levels

Simulated cell-scale water fluxes in the root cross-section anatomical network are illustrated in Fig. S2 (combining impermeable apoplastic barriers, high *k*_w_ and low *K*_PD_ and default boundary conditions from Tabs. 3 and 4, with air-filled intercellular spaces). Each panel puts the emphasis on the local variability of water flow paths at specific stages of apoplastic barriers development. Water fluxes in the apoplast are shown on horizontal surfaces (low to high water fluxes range from blue to red). Downward double-arrowheads point at parts of the apoplast containing hydrophobic substances or filled with air, thus blocking water capillary flow. Tangential cell walls also tend to have low fluxes (blue), simply because there is no water driving force oriented tangentially. Membrane fluxes are shown on vertical surfaces (red: water flows into the cell; blue: water is released into the apoplast through the plasma membrane). Fluxes through plasmodesmata (relative to the cell membrane surface) are shown as disks on vertical surfaces (red: water flows into the cell; blue: water is transferred to the neighbouring cell). Note that disks viewed from the top appear as ellipsoids, and may overlap. For each couple of neighbouring cells, a red disk necessarily pairs with a blue disk of the same intensity on the other side due to their relative perception of “inflow” and “outflow”. The observation angle determines which disk is visible in each pair.

Subplot (a) displays a close up on fluxes in the vicinity of an exodermis with a Casparian strip. The leftward arrowhead highlights that a significant fraction of water flows into exodermal cells through their outer tangential membranes (red vertical surface), while the rightward arrowhead shows water released on the other side, through the inner tangential membranes of the exodermis (blue vertical surface).

Subplot (b) focuses on another Casparian strip located at the endodermis, using the same colour scale. Tangential membranes (left and rightward arrowheads) display high fluxes due to the high radial conductivity of the young root segment, and to the small surface of the endodermis. A significant fraction of water flow that was not already released by inner endodermal membranes reaches the pericycle cells through plasmodesmata (red spots on vertical surfaces, see arrow). It is then released into the stelar apoplast through pericycle cell membranes (approximately 30% of the water follows this path as shown in Fig. 3c).

Subplot (c) displays a close up on fluxes in the vicinity of a partly suberized endodermis with two passage cells. Both inner and outer tangential walls are impermeable (see double arrowheads), which isolates most of endodermal membranes from apoplastic compartments. The leftward arrowhead shows that endodermal transmembrane flow still happens at passage cells sites (toward which apoplastic flow converges).

In contrast, in the fully suberized endodermis (no passage cells) of subplot (d), transmembrane flow takes exclusively place upstream in the cortex (leftward arrowhead) as no endodermal membrane is exposed to the apoplast. Arrows show plasmodesmata transferring water from cortical to endodermal cells, then to pericycle cells.

### Notes S3: Radial representation of water pathways

The radial representation aggregates the cell level water flow rates into pathways described on a cell layer basis, from the outer epidermal layer to the centre of the root. This visualisation is considered appropriate for the following reasons:

i. The relative contribution of symplastic and apoplastic water pathways to the total uptake varies widely across the root radius, particularly in the vicinity of apoplastic barriers where water flow in cell walls is interrupted [Enstone et al., 2002]. The proposed visualisation captures this variability.
ii. The radial view emphasizes the potential contribution of each cell layer in the root cross-section hydraulic network. For instance, the role of aquaporins expressed in specific cell layers at each stage of root anatomical development [Hachez et al., 2006; Hachez et al., 2012] may then be related to their functional relevance in the radial water pathway.
iii. Cells within the same layer bear similar hydraulic features and water status (at the condition that there is no significant asymmetry in soil water status). Hence, no significant trend was overlooked when aggregating cell scale information into a radial description.

The total flow rate at root surface *(Q*_s_, m^3^s^-1^) over the length of the root segment (200 microns) was used as reference 100% of flow rate in Fig. 3. As cells were arranged in layers connected in series from the root surface to the xylem vessels, and as no change in water content and no axial flow are considered (except for water leaving the cross-section through xylem vessels), the same amount of water flows across each cell layer from root surface to the first xylem vessel. When water reaches the xylem boundaries, where it feeds axial flow (see white boxes in Fig. 3), a decreasing fraction of *Q*_s_ flows toward inner layers.

Each cell layer being constituted of three hydraulic media assembled in series and parallel, the nature of flow across layers is intrinsically composite. In the radial representation, relative fractions of water flow in the apoplast and the symplast were quantified both within and between cell layers (see scheme in Fig. S3). Transmembrane flow (grey areas in Fig. S3b) is the only way to pass from apoplastic to symplastic pathways (respectively brown and blue areas in Figs. S3c and 3) and vice-versa. Hence, at the cell layer scale the transmembrane path does not add up to the first two pathways, but constitutes a transition between them. In consequence, the pathway typically referred to as “transcellular” [Steudle and Peterson, 1998] is here represented as repeated transitions between symplastic and apoplastic paths, respectively within and between cell layers (see particularly in Fig. 3c, f, i), rather than a separate third pathway.

The minimum set of information needed to quantify radial apoplastic and symplastic pathways within and between cell layers is the distribution of (i) water flow rates across plasma membranes and (ii) water flow rates exiting the root cross-section through xylem vessels (white boxes in Fig. 3).

#### Equations of water pathways at the cell layer level

In the following, *Q*_s_ (m^3^s^-1^) is the root surface water flow rate. Water flow rates across cell layers plasma membranes are noted *Q*_i,n_ and *Q*_o,n_, respectively into and out of the n-th cell layer (m^3^s^-1^), and flow rates leaving the system through xylem vessels are noted *Q*_x,n_ (m^3^s^-1^). The apoplastic and symplastic relative pathways (%) at the n-th cell layer transition (resp. *A*_T,n_ and *S*_T,n_, with *A*_T,0_ =100 and *S*_T,0_ = 0 at the root surface) were calculated as follows:
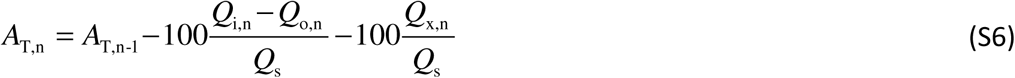

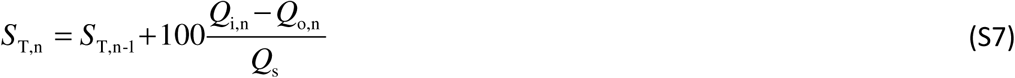

The relative apoplastic and symlastic flow within any n-th cell layer *A*_L,n_ and *S*_L,n_ (%, the first layer being the epidermis):
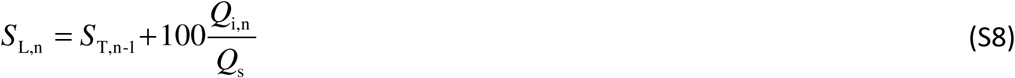

when *Q*_s_ is positive. Otherwise, the following equation applies:
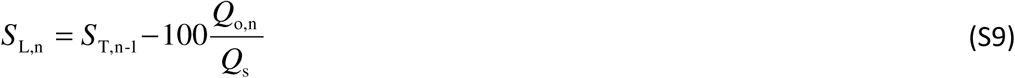

In case of significant asymmetry in the cross-section anatomy (more cell layers on a specific side), the radial layered view may be less appropriate as the layer at the centre of the cortex would form a croissant instead of a closed circle. However, absolute symplastic flow rates remain correct within the croissant, while the relative pathway would have to be locally adjusted accounting for the water flow rate bypassing the croissant. Within the range of xylem vessels and beyond, part (or all) of the flow leaves the cross-section axially, reducing the total radial flow accordingly (see white rectangle in Fig. 3). Note that for consistency between layers, all percentages in Figs. 3 and S3 are expressed relative to *Q*_s_.

### Notes S4: On the sensitivity analysis

The sensitivity (*L*_s_, %) of root *k*_r_ to specific local properties was quantified as the relative change of *k*_r_ divided by the relative change of that property in percent units:
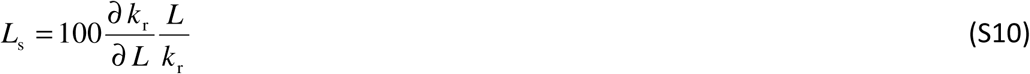

where *L* stands for the selected cell-scale hydraulic property (e.g., *L*_p_ or *K*_PD_).

Graphically, sensitivities correspond to the slopes of coloured line in Fig. 4. The sensitivity cannot exceed 100% by definition (1:1 ratio) for the following reasons:

The sensitivity of *k*_r_ to a type of conductance (*K*_1_) is 100% in a network composed of that single conductance; adding a different type of conductance (*K*_2_) in parallel decreases the sensitivity down to 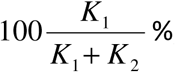, and the sum of both sensitivities equals 100%.

Similarly, adding *K*_2_ in series decreases the sensitivity of *K*_1_ down to 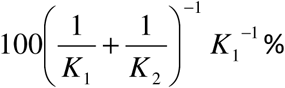, and the sum of both sensitivities equals 100%.

As the hydraulic network is constructed by progressively adding conductances in series and parallel, the sensitivities of *k*_r_ to complementary elements of the hydraulic network must be comprised between 0 and 100%, and sum up to 100%.

**Figure S1.**
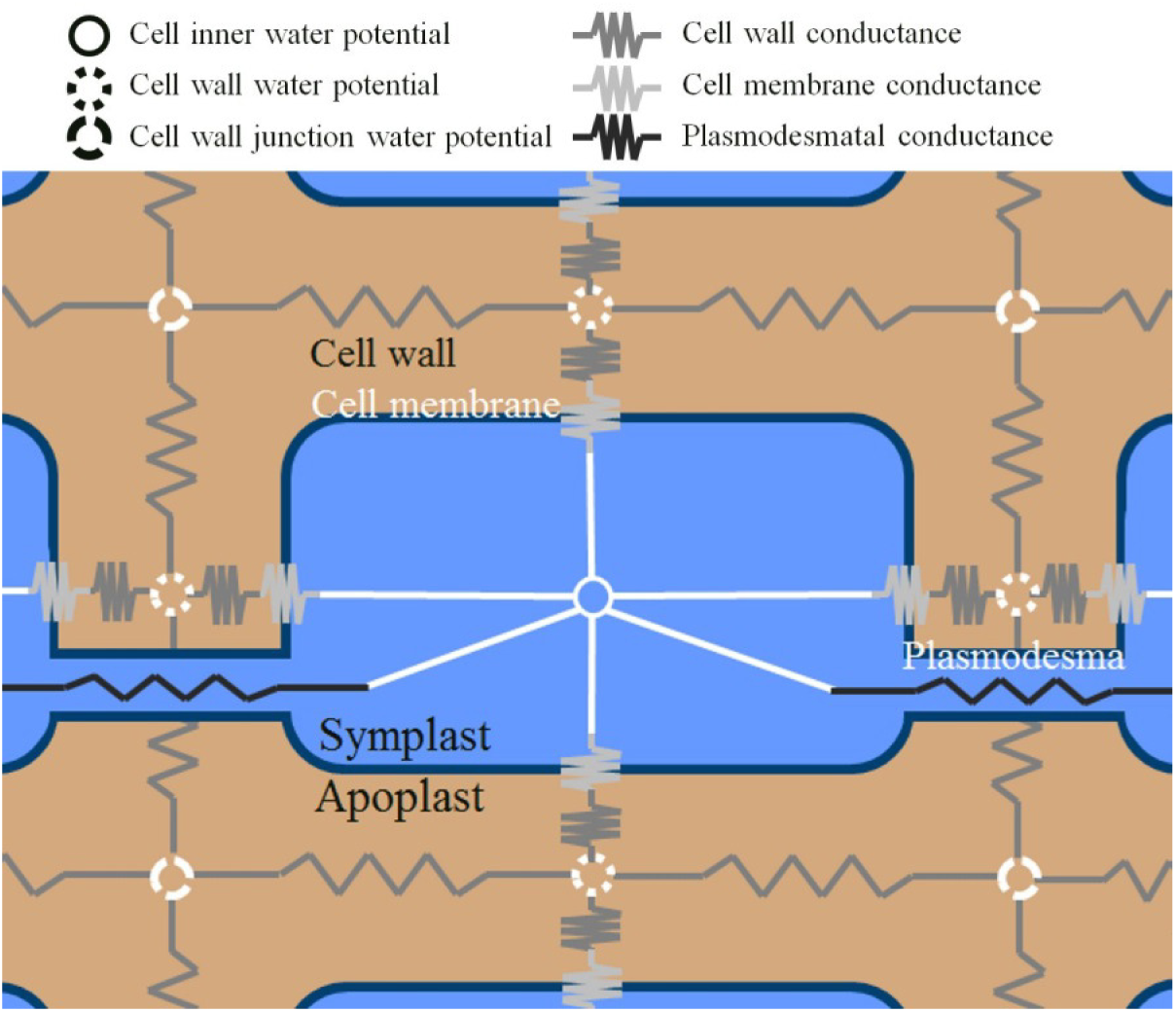
Scheme of the cell-level hydraulic network. Each cell protoplast (blue) is discretised into a single node connected to the surrounding apoplast (brown) by transmembrane hydraulic conductances (light grey) and a short wall hydraulic conductance (dark grey) to reach the centre of the space between protoplasts. Cell wall junction nodes connect cell wall nodes to each other with longer wall hydraulic conductances (dark grey). Protoplasts of neighbouring cells are also directly connected by plasmodesmatal hydraulic conductances (black).

**Figure S2.**
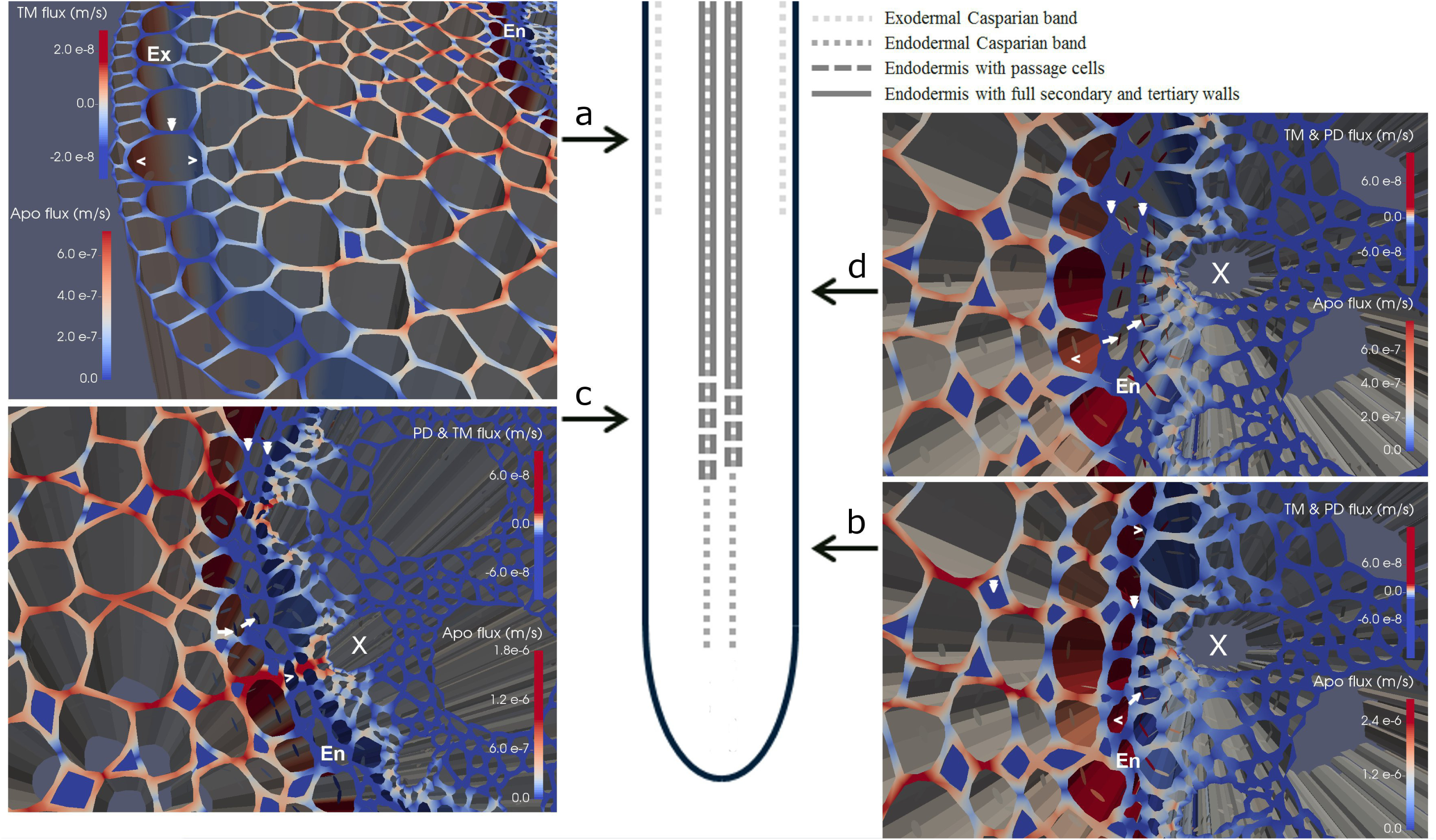
Visualization of water fluxes at different stages of apoplastic barrier development for impermeable apoplastic barriers, high *k*_W_, low *K*_PD_ and default boundary conditions (see Tabs. 3 and 4). Apoplastic fluxes are displayed on horizontal surfaces, high and low fluxes appear in red and blue, respectively. Cell membrane fluxes are displayed as vertical surfaces, positive (red) into the symplast, and negative (blue) leaving the symplast. Plasmodesmatal fluxes are displayed as disks on vertical surfaces, positive (red) into the cell and negative (blue) leaving the cell. Ex, Exodermis; En, Endodermis; X, Xylem. (a) Exodermal Casparian strip; (b) Endodermal Casparian strip; (c) Endodermal passage cells; (d) Fully suberized endodermis. Downward double arrowheads point at hydrophobic cell walls and air-filled intercellular spaces. Arrowheads point at cell plasma membranes. Arrows point at plasmodesmata.

**Figure S3.**
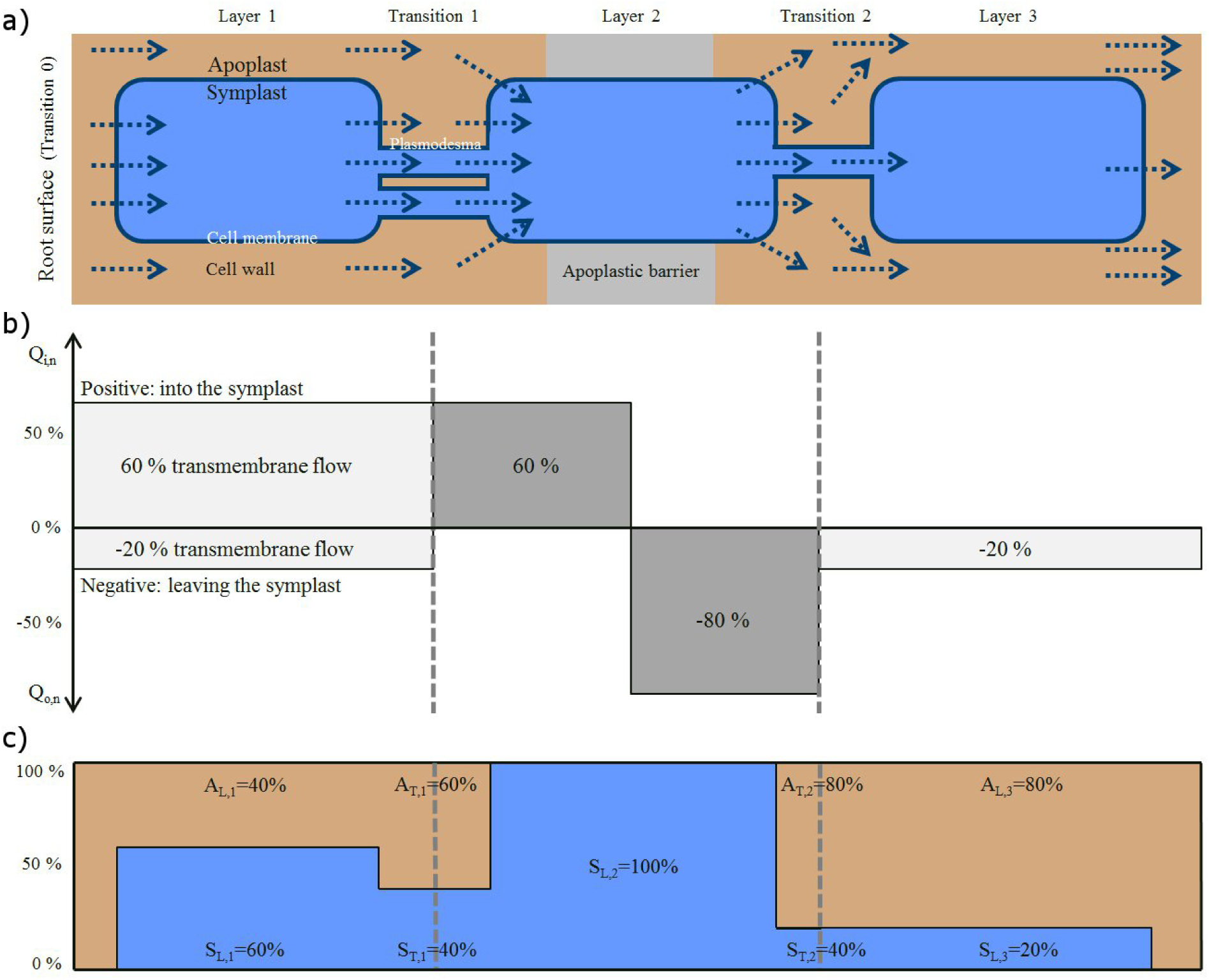
Scheme of the method used to calculate radial water pathways from local transmembrane flow rates. (a) Simple triple cell network. Each dotted blue arrow represents one unit of water flow. Overall the network is crossed by 5 units of water flow, from root surface (left) to the stelar apoplast (right). Due to mass conservation, the same total flow rate enters and exits each cell layer (no xylem vessels represented here). (b) Transmembrane flow rates of the same layer are summed up to obtain each layer’s *Q*_i,n_ (positive, apoplast to symplast) and *Q*_o,n_ (negative, symplast to apoplast). Note that transmembrane flow rates on both sides of an apoplastic barrier are represented separately and appear in dark grey. When the root takes up water, the sum of terms on the left (resp. right) of the apoplastic barrier sum up to 100% (resp. -100%) (c) Pathways are calculated within cell layers and at their transitions based on transmembrane flow rates, which quantitatively mark pathway changes (from apoplast to symplast, or vice versa). Within cell layers that bear an apoplastic barrier, the symplastic pathway *S*_L_ reaches 100%.

